# Type IV pilus shapes a ‘bubble-jet’ pattern opposing spatial intermixing of two interacting bacterial populations

**DOI:** 10.1101/2021.10.17.464652

**Authors:** Miaoxiao Wang, Xiaoli Chen, Yinyin Ma, Yue-Qin Tang, David R Johnson, Yong Nie, Xiao-Lei Wu

## Abstract

Microbes are social organisms that commonly live in sessile biofilms. Spatial patterns of populations within biofilms can be an important determinant of community-level properties. The best-studied characteristics of spatial patterns is spatial intermixing of different populations. The specific levels of spatial intermixing critically contribute to how the dynamics and functioning of such communities are governed. However, the precise factors that determine spatial patterns and intermixing remain unclear. Here, we investigated the spatial patterning and intermixing of an engineered synthetic consortium composed of two *Pseudomonas stutzeri* strains that degrade salicylate via metabolic cross-feeding. We found that the consortium self-organizes across space to form a previously unreported spatial pattern (referred to here as a ‘bubble-jet’ pattern) that exhibits a low level of intermixing. Interestingly, when the genes encoding for type IV pili were deleted from both strains, a highly intermixed spatial pattern developed and increased the productivity of the entire community. The intermixed pattern was maintained in a robust manner across a wide range of initial ratios between the two strains. Our findings show that the type IV pilus plays a role in mitigating spatial intermixing of different populations in surface-attached microbial communities, with consequences for governing community-level properties. These insights provide tangible clues for the engineering of synthetic microbial systems that perform highly in spatially structured environments.

**Importance:** When growing on surfaces, multi-species microbial communities form biofilms that exhibit intriguing spatial patterns. These patterns can significantly affect the overall properties of the community, such as enabling otherwise impermissible metabolic functions to occur, as well as driving the evolutionary and ecological processes acting on communities. The development of these patterns is affected by several drivers, including cell-cell interactions, nutrient levels, density of founding cells and surface properties. The type IV pilus is commonly found to mediate surface-associated behaviors of microorganism, but its role on pattern formation within microbial communities is unclear. Here we report that in a cross-feeding consortium, the type IV pilus affects the spatial intermixing of interacting populations involved in pattern formation, and ultimately influences overall community productivity and robustness. This novel insight assists our understanding of the ecological processes of surface-attached microbial communities and suggests a potential strategy to engineer high-performance synthetic microbial communities.

---

In addition to the planktonic lifestyle, microorganisms also form intricate multispecies communities on surfaces (1, 2). These surface-attached communities play important roles in ecosystem processes (3), pollutant removal (4) and human health (5). Biofilms are spatially well-organized, with different populations interacting with each other, arranging themselves non-randomly across space, and ultimately developing well-organized spatial patterns (referred to as ‘spatial self-organization’ hereafter (6, 7)). Spatial patterns of a community reflect the distribution of different populations across the habitat, and this distribution profoundly influences the interactions that occur among these populations. For example, spatial mixing of different interacting cells (referred to as a ‘mixed pattern’ thereafter) benefits their metabolic exchanges (8) and alleviates antibiotic stress (9). On the other hand, spatial demixing (referred to as ‘segregated pattern’ thereafter) stabilizes intransitive interactions among antibiotic-producing, -sensitive and -resistant species (10), and also protects cells from contact-dependent killing by its competitors (11). These outcomes can eventually be magnified to determine community-level properties, such as overall community productivity (8), resistance to invaders (12, 13), and robustness to initial conditions (7, 14).

Recently, several abiotic and biotic factors have been reported to influence the intermixing level of spatial patterns. For instance, genetic surfing during range expansion generally demixes populations during growth, resulting in a segregated pattern (15). However, the degrees of intermixing can be increased by increasing the nutrient levels (16), increasing the density of founding cells (17), the presence of physical objects (18) and promoting metabolic interactions (19, 20). As a result, the community will self-organize into a more intermixed pattern. In addition, several studies reported cell appendages, such as Type IV pilus, play an essential role in microbial colonization on surface (21). On the one hand, Type IV pilus mediates twitching motility of bacterial cells (21). On the other hand, it also contributes to stabilizing interactions between cells and the abiotic surface (22), as well as facilitating cell-to-cell adhesion (23, 24), which is required for aggregate formation. Nevertheless, whether this functional cell appendage impacts the intermixing level of the formed spatial patterns still remains to be elucidated.

Here, we explored a selection of factors potentially affecting spatial self-organization of a well-defined two-strain consortium using the strains *Pseudomonas stutzeri* AN0010 and *P. stutzeri* AN0001. While strain AN0010 degrades salicylate into the intermediate catechol, strain AN0001 further degrades catechol to pyruvate and acetyl-CoA (Figure 1A). We previously showed that when paired, these two strains engage in a cross-feeding interaction in the presence of salicylate (25).

**Figure 1.**
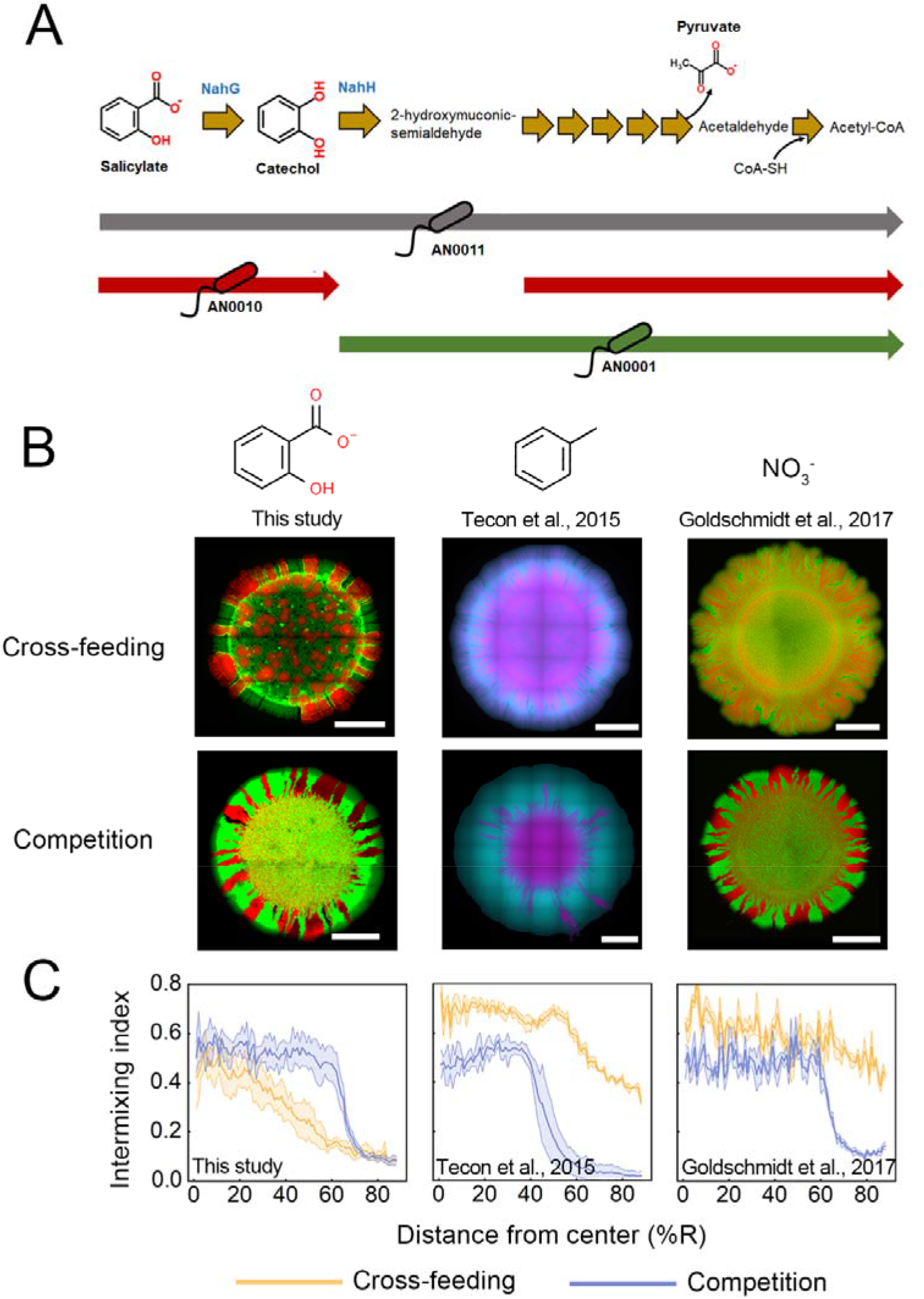
The salicylate-degrading community self-organized into a ‘bubble’ jet pattern characterized by lower intermixing level. (A) Pathway of salicylate degradation in strain *P. stutzeri* AN0011, as well as partial pathways carried out by strain AN0010 and strain AN0001. (B) Representative colony patterns developed by our salicylate-degrading community, as well as patterns developed by the similar cross-feeding communities previously constructed. Overlay fluorescence images of these patterns are shown. For the patterns formed by our salicylate-degrading community (left), strain AN0010 was tagged with mCherry (shown as pseudo-color Red), while strain AN0001 were labeled with eGFP (shown as pseudo-color Green). Shown in the middle panels are the patterns developed by a toluene-degrading cross-feeding community built by Tecon (14) *et al*. (2015), in which strain *P. putida* PpF4 (tagged with eGFP and shown as cyan) degrades toluene to 3-methylcatechol, before it being transformed to acetate and pyruvate by strain *P. putida* PpF107 (tagged with mCherry and shown in magenta). Shown in the right panels are the patterns developed by a denitrification cross-feeding community built by Goldschmidt (26) *et al*. (2017), in which strain *P. stutzeri* A1603 (tagged with eCFP and shown as green) converts NO_3_^-^ to NO_2_^-^, and strain *P. stutzeri* A1602 (tagged with mCherry and shown as red) further reduces NO_2_^-^ to N_2_. For all three synthetic communities, pattern formation assays in ‘Cross-feeding’ scenario (supplying salicylate, toluene or NO_3_^-^ in the media, respectively) and ‘Competition’ scenario (supplying pyruvate, benzoate as the sole carbon source, or grow two complete NO_3_^-^ degraders together on LB agar surface). The scale bar in each image corresponds to 1 mm. (C) Analysis of intermixing indexes of these patterns. Higher values indicate higher levels of local spatial intermixing of the two strains. These values were assessed through image analysis following the protocol modified from a previous study (26) (see Supplementary Information 1.5 for details). To obtain the patterns formed by our salicylate-degrading community, six experimental replicates were performed. Images of the patterns formed by the toluene-degrading community were published (14), courtesy of Dr. Robin Tecon. In addition, the microscopic images of the patterns formed by denitrification community were obtained by performing three replicated pattern formation assays following a protocol previously reported (26).

To investigate the spatial self-organization of this community, we cultured the community on an agarose surface using salicylate as the sole carbon source. Based on findings from previous studies, we expected our cross-feeding community to self-organize into a highly mixed pattern (14, 19, 20, 26). Surprisingly, we observed instead a segregated pattern, where the cells of strain AN0010 formed bubble-like structures inside the colony, with cells of strain AN0001 subsequently surrounding these bubbles (Figure 1B; Figure S1A). During range expansion, cells of strain AN0010 expanded from these ‘bubble’ structures, similar to the cells being ‘jetted’ outside of the bubbles to form the expanding sectors. We therefore refer to this previously undescribed spatial pattern as the ‘bubble-jet’ pattern. Our analysis using three-dimensional confocal microscopy showed that cells of strain AN0010 assembled in the ‘bubble’ structure, exhibiting a ‘bowl’-like geometrical morphology (Figure S1B). Further analysis of the fluorescence intensities of the two genotypes showed that cells of strain AN0001 were mostly distributed around the bubble formed by strain AN0010 (Figure S1C), suggesting that the metabolic interaction between the two populations still necessitated that the two populations were located in close spatial proximity.

To quantitatively test whether the levels of mixing between the two populations within the ‘bubble-jet’ pattern are lower than the previously reported patterns developed by other cross-feeding consortia, we calculated the intermixing indexes of all these patterns (26) (see Supplementary Information S1.5 for details). We found that the intermixing levels of the ‘bubble-jet’ pattern are significantly lower than those generated by cross-feeding consortia performing toluene degradation (14) (Mann–Whitney test, *p* = 1.07e-23) and denitrification (26) (Mann–Whitney test, *p* = 2.64e-28; Figure 1C). In addition, previous studies reported that spatial intermixing in those cross-feeding communities was reduced when the strains become metabolically independent (14, 19, 20, 26). We thus set out to test whether eliminating metabolic cross-feeding in our synthetic community also reduce the intermixing levels of the ‘bubble-jet’ pattern. We cultured our synthetic community on an agar surface using pyruvate as the sole carbon source. Pyruvate constitutes one of the final products of the salicylate degradation pathway, and is directly utilized by both strains for growth (Figure 1A). When the two strains directly competed for this limited resource, a clear segregated pattern formed (Figure 1B). However, our analysis of this pattern suggested that the ‘bubble-jet’ pattern formed in the ‘cross-feeding’ scenario is even less mixed than the pattern formed in the ‘competition’ scenario (Figure 1C; Mann–Whitney test, *p* = 4.61e-9), in disagreement with previous observations (14, 26) (Figure 1B-C). Moreover, we also tested whether the ‘bubble-jet’ pattern is observed regardless of the magnitude of known key drivers, for example, nutrient levels (16) and density of founder cells (17). These investigations indicated that although these factors quantitatively affected the detailed morphology of the pattern, alteration of these factors failed to qualitatively affect the development of the ‘bubble-jet’ pattern (Figure S2-S3).

It has previously been reported that cell appendages, such as type IV pilus and flagellum, are critically involved in the formation of *Pseudomonas* biofilms (22, 27). To investigate whether these cell appendages are also involved in the formation of our ‘bubble-jet’ pattern, we introduced loss-of-function deletions in genes encoding key proteins involved in pilus and flagellum assembly of both strains (Figure S4). Interestingly, while the deactivation of flagellar genes did not change the development of the ‘bubble-jet pattern’. In comparison, deactivation of the type IV genes encoding pilus caused the ‘bubble-jet’ pattern to disappear and significantly increased the spatial intermixing of the two interacting populations in the developed pattern (Figure 2A-B; Mann–Whitney test, *p* = 3.96e-27). The mixed pattern that formed also better resembled the ones that developed in previous similar studies ((14, 26); Figure 2 A-B; Mann–Whitney test of the intermixing indexes: *p* = 0.026 compared with the pattern formed by the toluene-degrading community, and *p* = 0.017 compared with that of the denitrification community), and showed higher intermixing compared to the pattern formed by the same community in the ‘competition’ scenario (Mann–Whitney test, *p* = 6.06e-17). To test whether the disappearance of the ‘bubble-jet’ pattern only requires the pili mutation in a single strain, we examined the patterns formed by mixing the Δpili mutant of one strain to the wild-type of the other strain. We found that the ‘bubble-jet’ pattern completely disappeared once the pili of strain AN0010 were knocked out (Figure S5A). Moreover, the size of ‘bubbles’ significantly reduced when the pili mutant of AN0001 was mixed with the strain AN0010 (Figure S5D-E). Together, these results strongly suggest that the presence of type IV pilus is a determining factor in the formation of the to ‘bubble-jet’ pattern and controls the intermixing level of the spatial pattern.

**Figure 2.**
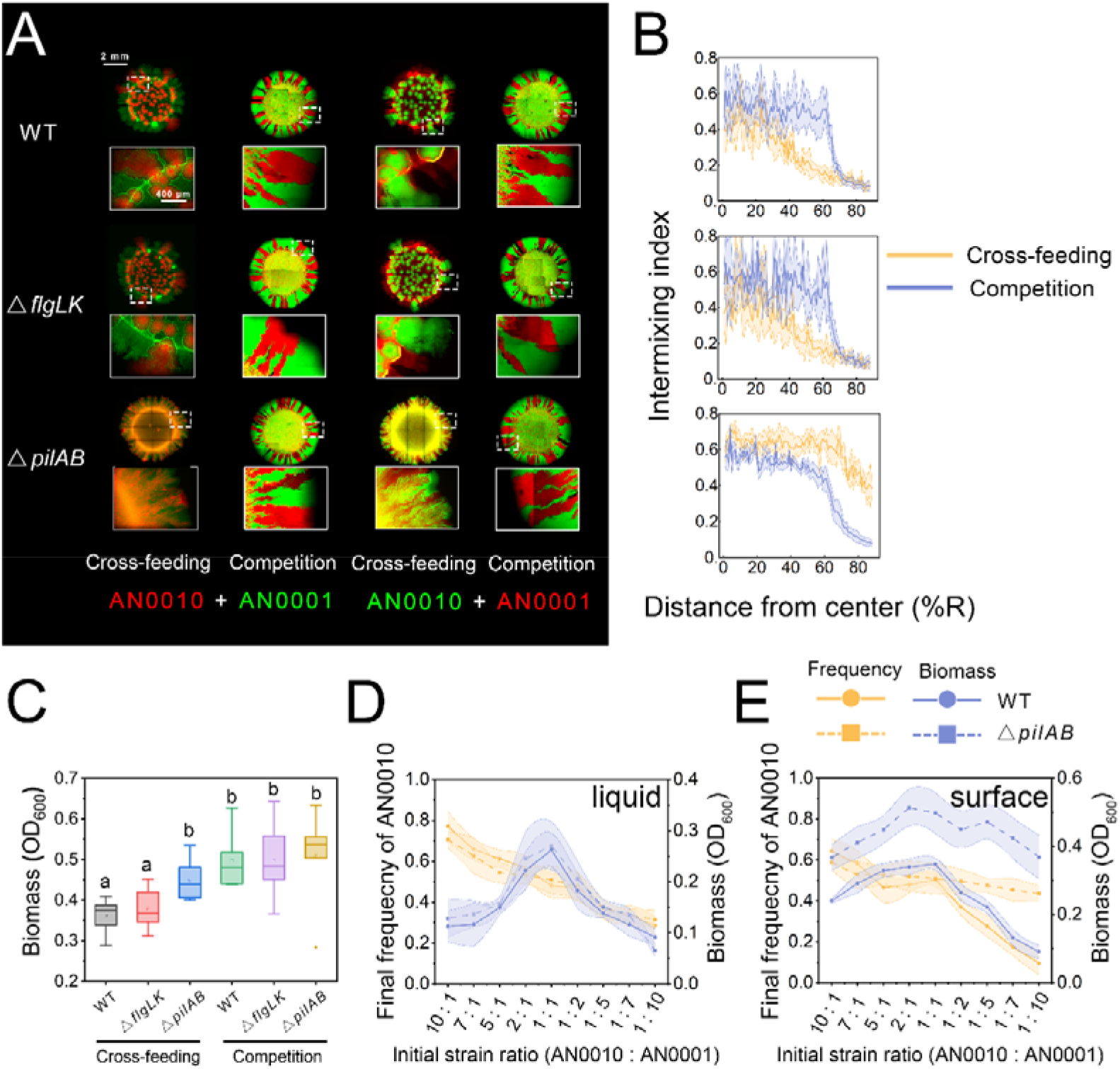
Type IV pili are required for formation of the ‘bubble’ structures, while flagella are dispensable. (A) Images showing that the colony patterns formed by co-culturing the wild type strain of AN0010 and AN0001 (top), their flagellum-mutant strains (middle), as well as their pili-mutant strains (bottom) on agarose surface. Patterns obtained in both ‘Cross-feeding’ scenario (supplying salicylate as the sole carbon source) and ‘Competition’ scenario (supplying pyruvate as the sole carbon source) were shown. Alternative fluorescence labelling was used to eliminate potential effects caused by expression of different fluorescent proteins. Typical morphology of colony edges (five-time zoom). Images were obtained after 120-h incubation. (B) Analyses of intermixing index of these patterns. (C) Biomass analysis of colonies. Lower-case letters indicate significant differences between these conditions at 0.05 level (unpaired, two-tailed, Student’s t-test). (D-E) The alterations of the salicylate-degrading community composed of the wild type strains (solid lines), as well as the community composed of the pili-mutant (dashed lines), against the initial strain ratios between the two populations. We cultured these communities initiated with nine different strain ratios. After incubation (144-h for liquid cultivation and 120-h for cultivation on agarose surface), final community structures (yellow line) and biomass (blue line) were analyzed. In liquid cultivation, both communities exhibited similar alterations in community structures and the biomass against different strain ratios (D). However, when the communities grew in agarose surface, for the community composed of pili-mutant strains, the final ratio of the two strains converged to similar values (≈ 1:1) regardless of initial strain ratio (E). In comparison, the final strain ratio for the community composed of wild type strains exhibited larger variations. Furthermore, the initial strain ratio also showed smaller effects on the final productivity (biomass) of the pili-mutant community than that of the wild-type community. These results strongly suggested that the community formed by pili mutants is more robust against any fluctuations in the initial conditions. For each initial strain ratio, six replicated experiments were performed. See Table S1 for quantitative comparison of variations in different conditions.

To investigate whether increased intermixing following removal of type IV pili influences community-level properties, we compared the biomass of the colonies developed by the communities composed of wild type strains and the Δpili mutant strains. Intriguingly, although the two communities grew similarly in liquid culture (25), colonies developed by the Δpili mutants produced more biomass than the wild type strains (Figure 2C, unpaired two-tailed Student’s t-test, *p* = 0.011; Figure S5C). This result suggested that Δpili mutants of the two strains interact better with each other than the wild type strains, possibly due to the spatial proximity to each other in the more mixed pattern, leading to increased productivity at the community level. We next tested the robustness of community structure and productivity to the initial strain ratio, which was previously reported to constitute a key feature of the cross-feeding community (14, 19). Despite the fact that the two communities exhibited similar robustness to the initial strain ratio in liquid cultivation (Figure 2D; see Table S1 for variation comparisons), we found that on agarose surface, the community composed of pili mutants was more robust to the initial conditions than the community that formed the ‘bubble-jet pattern’ (Figure 2E; Table S1). Together, these results implied that removing the pili increased the overall productivity, and renders the community for robust against varied initial ratios between the two strains.

Several studies based on microscopic investigation directly observed that *Pseudomonas* cells harboring Type IV pilus could migrate along the surface and gather together by recruiting adjacent and homologous cells, resulting in the formation of multicellular aggregates (22, 28). Therefore, we hypothesize that the observed ‘bubble’-like structures in our study may be derived from pilus-mediated formation of cell aggregates. However, this hypothesis requires further testing on the single-cell level.

Here, we found that a cross-feeding community self-organized into a previously unknown ‘bubble-jet’ pattern in the presence of pili structures, which opposes spatial mixing of the different populations involved. Our findings also demonstrated that the reduced spatial mixing is associated with a decrease in community productivity and robustness. In conclusion, our findings add critical insights to our current understanding of the ecology of surface-attached microbial communities. In addition, our results strongly suggest a potential strategy to engineer artificial communities with optimized spatial patterns – engineer the pili of the interacting strains to modulate interspecific distances in a surface-attached community, and thus we can promote its performance.

## Acknowledgments

We wish to thank Dr. Min Lin (Chinese Academy of Agricultural Sciences, Beijing, P.R. China) for providing plasmid pK18mobsacB and pRK2013, used for genetic engineering in this work; Professor Ping Xu (Shanghai Jiao Tong University, Shanghai, P.R. China) for supplying plasmid pMMPc-Gm, used for fluorescence labeling in this study; Professor Dani Or (ETH Zurich, Zurich, Switzerland) and Dr. Robin Tecon (NCCR Microbiomes and University of Lausanne) for providing the source images of their previous study; Professor Martin Ackermann (ETH Zurich, Zurich, Switzerland) for constructive inputs on the design of this study; Dr. T. Juelich (UCAS, Beijing) for linguistic assistance during the preparation of this manuscript. This work was supported by National Key R&D Program of China (2018YFA0902100 and 2018YFA0902103), and National Natural Science Foundation of China (32130004, 91951204, 31770120, and 31770118).

