## Supplementary Information for "Type IV pilus shapes a ‘bubble-jet’ pattern opposing spatial intermixing of two interacting bacterial populations"

### S1 Materials and Methods

#### *S1.1 Bacterial strains*

The two *Pseudomonas stutzeri* strains involved in our synthetic salicylate-degrading community were engineered in our previous work (1). Their biological traits were summarized in Figure 1A. Briefly, strain *P. stutzeri* AN0011 can degrade salicylate autonomously, because it contains all the genes encoding the corresponding enzymes that are located in an operon induced by IPTG. To generate *P. stutzeri* AN0010 and *P. stutzeri* AN0001, *nahG* (encoding a salicylate 1-hydroxylase) or *nahH* (encoding a catechol 2,3-dioxygenase) gene of *P. stutzeri* AN0011 was knocked out, respectively. Thus, strain AN0010 is only capable of transforming salicylate to catechol, while strain AN0001 that can only degrade catechol. Furthermore, catechol cannot be directly used as the carbon source by both strains, while the final products of this pathway, i.e., pyruvate and Acetyl-CoA (Figure 1A) represent the direct carbon sources for the two strains in this community. Therefore, when co-cultured using salicylate as the sole carbon source, they exchange intermediate catechol and final product pyruvate, exhibiting a cross-feeding interactions.

To generate the pilus mutants or flagellum mutants, *pilA* and *pilB* genes, or *flgL* and *flgK* genes, were simultaneously removed from the host strains, respectively. The genetic manipulation was implemented by allele exchange using the suicide plasmid pK18mobsacB (2, 3). The constructed strains were validated by PCR and DNA sequencing. Furthermore, agar-based 'stab' assays were performed to validate the pilus mutants (Figure S4B), while 'swimming' assays were performed to validate flagellum mutants (Figure S4C), both following the standard protocol (4). To label the strains with fluorescence, mCherry or eGFP was cloned into a constitutive vector, pMMPc-Gm (5), and delivered to the host cells via triparental filter mating (3).

*S1.2 Colony pattern formation assays*

Minimum medium (1.5% agarose, Takara, Dalian, China), supplemented with 2 mM IPTG, 50 μg/mL gentamicin, and supplying salicylate or pyruvate (10 Cmmol/L unless otherwise indicated) as sole carbon source, was used in these studies. To prepare the culture plate, five milliliters of this medium was poured in a Petri dish (60 mm in diameter) and left on the bench overnight before inoculation. To prepare the inoculum, *P. stutzeri* strains were first grown at 30˚C RB liquid medium (Yeast extract 10 g/L, beef extract 6 g/L, peptone 10 g/L, ammonium sulfate 5 g/L) by shaking at 220 rpm, supplemented with 50 μg/mL gentamicin. The cells were then washed by the minimum medium (6) for twice to make an inoculum. Inocula of two strains were then concentrated to an Optical Density (OD, measured at 600 nm) of 1.0, and mixed at a 1:1 ratio. For each colony, 1 μl of the inoculum was spotted on the prepared plate, after which the agar plate was allowed to dry for 10 min. After the inoculum dried on the plate, the plates were incubated at 30 ˚C for 120 h.

In addition, to test the effect of substrate concentration (Figure S2), final concentration of salicylate in the medium was varied from 2.5 Cmmol to 20 Cmmol. To test the effects of cell density (Figure S3), the Inocula were concentrated to ODs varied from 0.05 to 10.0. To test the robustness of the community to initial strain ratio (Figure 2D), the two strains were initially mixed at a ratio ranging from 1 : 10 to 10 : 1.

*S1.3 Microscopy imaging*

Colony patterns were imaged under 5× objective using a Leica DM6000B fluorescence microscope (Leica Corporation, Wetzlar, Germany) equipped with a LED fluorescence illuminator (Leica Corporation). Images were sequentially recorded with a DFC360 FX camera (Leica Corporation) using a GFP filter cube for eGFP (exciter: 475/40; emitter: 525/50: beamsplitter: 495) and a TX2 filter cube for mCherry (exciter: 560/40; emitter: 645/75: beamsplitter: 595). Tile scan function of Leica LAS X acquisition software (Leica Microsystems) were applied to assemble the full view of a colony from multiple fields, and composite images were also created by this software.

*S1.4 Confocal imaging*

To investigate the 3D structure of the 'bubble'-like structure (Figure S1B), typical bubble areas were imaged under 10× objective using an Andor Revolution XD laser confocal fluorescence microscope (Andor, Oxford, UK) associated with ANDOR Zyla sCOMS camera. Individual color channels were acquired using the FLIC and TxRed filters in addition to a bright-field channel. The images were assembled and virtualized in Image J software (version 1.53c). In addition, 'Surface Plot' function of Image J software was applied to analyze the relative intensity of fluorescence across the bubble areas (Figure S1C).

*S1.5 Imaging analysis.*

Wolfram Mathematica (version 12.4) was used to process and analyze images. Firstly, the assembled eGFP and mCherry images were separately exported as grayscale tiff files, and then resized into images with 2000 pixels wide (using *ImageResize* function) to reduce computation time. Then *ColorQuantize* function was used to give an approximation to the image by quantizing it to distinct colors, subsequently transform the images into binarized data using *ImageData* function. Using these binarized matrix data as the input, we quantified the intermixing level of different patterns by evaluating their intermixing indexes following a protocol modified from previous study (7). The intermixing index at a given radius ($r$) was defined as the number of intersections between different populations ( $N_r$ ) normalized by the circumference of the corresponding circle, as follow:

$$I_r = \frac{N_r}{2\pi r}$$

Therefore, $I_r$ estimates the intermixing level of the two populations (colored pixels) at a distance $r$ from the colony center. In particular, $I_r$ equaling 0 means that all pixels around the circumference only contain one or the other strain; $I_r$ equaling 1 means that all pixels around the circumference contain equal concentrations of both colors and they are alternatively distributed around the circumference with the highest number of intersections. Intermixing indexes are shown as a function of distance ($r$). Generally, intermixing is relatively constant in the inoculation zone, then decreases in the expansion zone, except when the 'bubble-jet' pattern was developed (e.g., Figure 1).

We quantified the morphological changes of the 'bubbles' by performing a segmentation imaging analysis of the 'bubble' area (Figure S2B). Image with the color channel related to AN0010 was used. We first cropped the images to focus on the inside area containing 'bubbles'. Then the 'bubbles' were segmented by the watershed algorithm using the *WatershedComponents* function of the *Wolfram Mathematica*, followed by obtained the statistic data of the number and size of these structures by using *ComponentMeasurements* function. The area size in pixels were finally transformed to be the real size (in $mm^2$) by multiply the scale bar, and virtualized by *Colorize* function.

All these calculations were performed in custom *Wolfram Mathematica* scripts. The source codes used are available on Github:

https://github.com/RoyWang1991/MDOLcode/tree/master/MDOL-spatial.

*S1.6 Measurement of biomass and final strain ratio*

To estimate the biomass and community composition of a colony, spot was collected by an inoculation loop from the plate after 120-h's culture, and bacterial cells were resuspended in 100 μL minimum medium and vortexed for 10 min to destruct the biofilms. After that, the OD was measured to estimate the total biomass, and fluorescence intensities of eGFP and mCherry were measured to estimate the growth of each population. The final strain ratio was calculated by a method previously described (1, 8). Briefly, cultures of each populations were grown to mid-log phase at 30˚C (OD: ~0.3), diluted two-fold for eleven times, and the dilutions measured for their OD and fluorescence. Correlations between OD and fluorescence were then determined using the basic method defined in the LinerModelFit function of *Wolfram Mathematica* software (version 12.4). Eventually, fluorescence values were transformed to the OD-estimated biomass to assess the growth of each population, and relative fraction was then calculated. These measurements, as well as the related measurements described below, were performed using a microplate reader (Molecular Devices, Sunnyvale,

America).

*1.7 Liquid cultivation of the P. stutzeri strains*

Firstly, inocula is prepared by the same way as described in section 1.6. For co-culture experiments, inocula of two strains the were concentrated to an Optical density (OD, measured at 600 nm) of 5.0, and mixed at different initial ratios, and then inoculated to 96-well plates that contains 120 μL fresh minimum medium (starting OD: 0.05), supplemented with 2 mM IPTG, 50 μg/mL gentamicin, as well as salicylate as the sole carbon source (final concentration 10 C-mmol). During the cultivation, OD was measured to estimate the total biomass, and fluorescence intensity was measured to estimate the growth of each population. Calculation of the relative frequency followed the same method described in the section 1.6.

*1.8 Statistical analyses*

Unless indicated otherwise, the number of replicates is six for each experiment. Mann–Whitney test was used for statistically comparative the intermixing level of different patterns, while unpaired, two-tailed, Student's t-test was applied for other comparative statistical analyses. These analyses were performed in *Wolfram Mathematica* (version 12.4).

### S2 Supplementary Figures

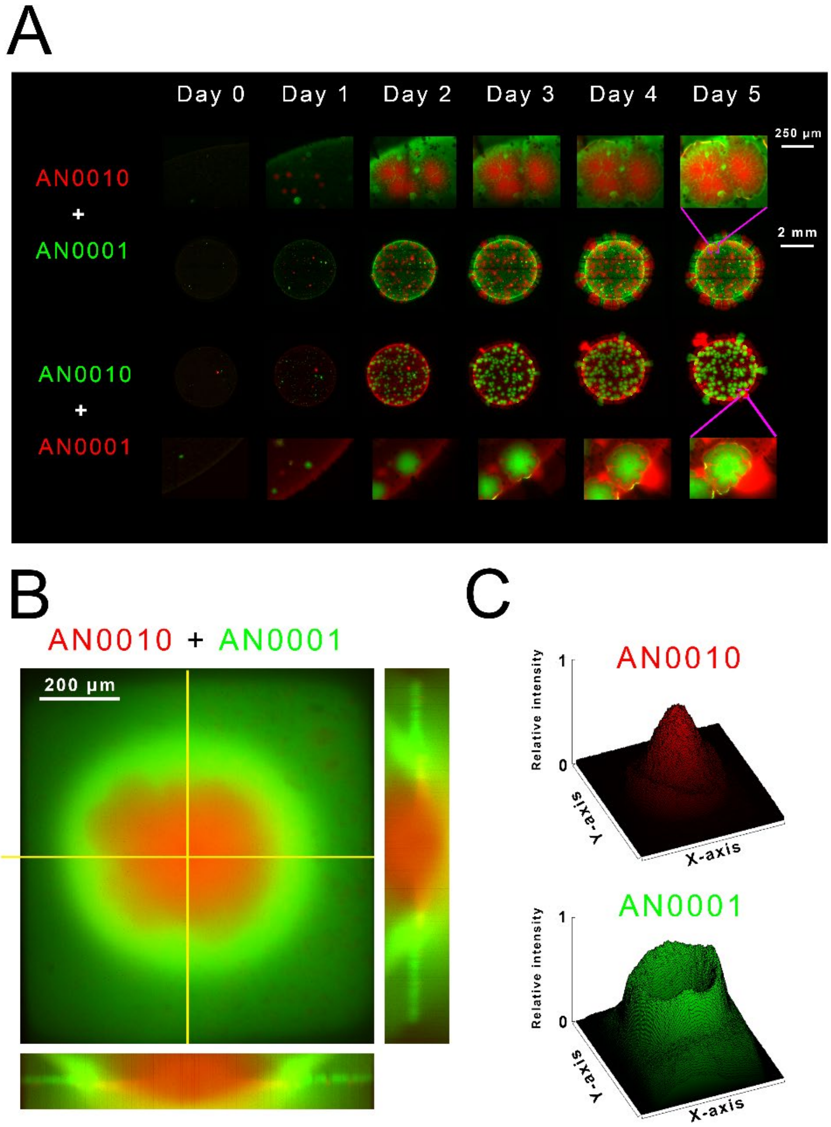


**Figure S1** Characterization of the spatial pattern formed by our synthetic salicylate-degrading community. (A) Images show the colony growth dynamics of the community when supplying salicylate as the sole carbon source. The growth of typical 'bubble' areas is zoomed in. (B) Confocal imaging shows the three-dimensional structure of a typical 'bubble' area. (C) Analysis of the relative fluorescence intensity of the image showed in (B), suggesting the distribution of the two populations in this area.

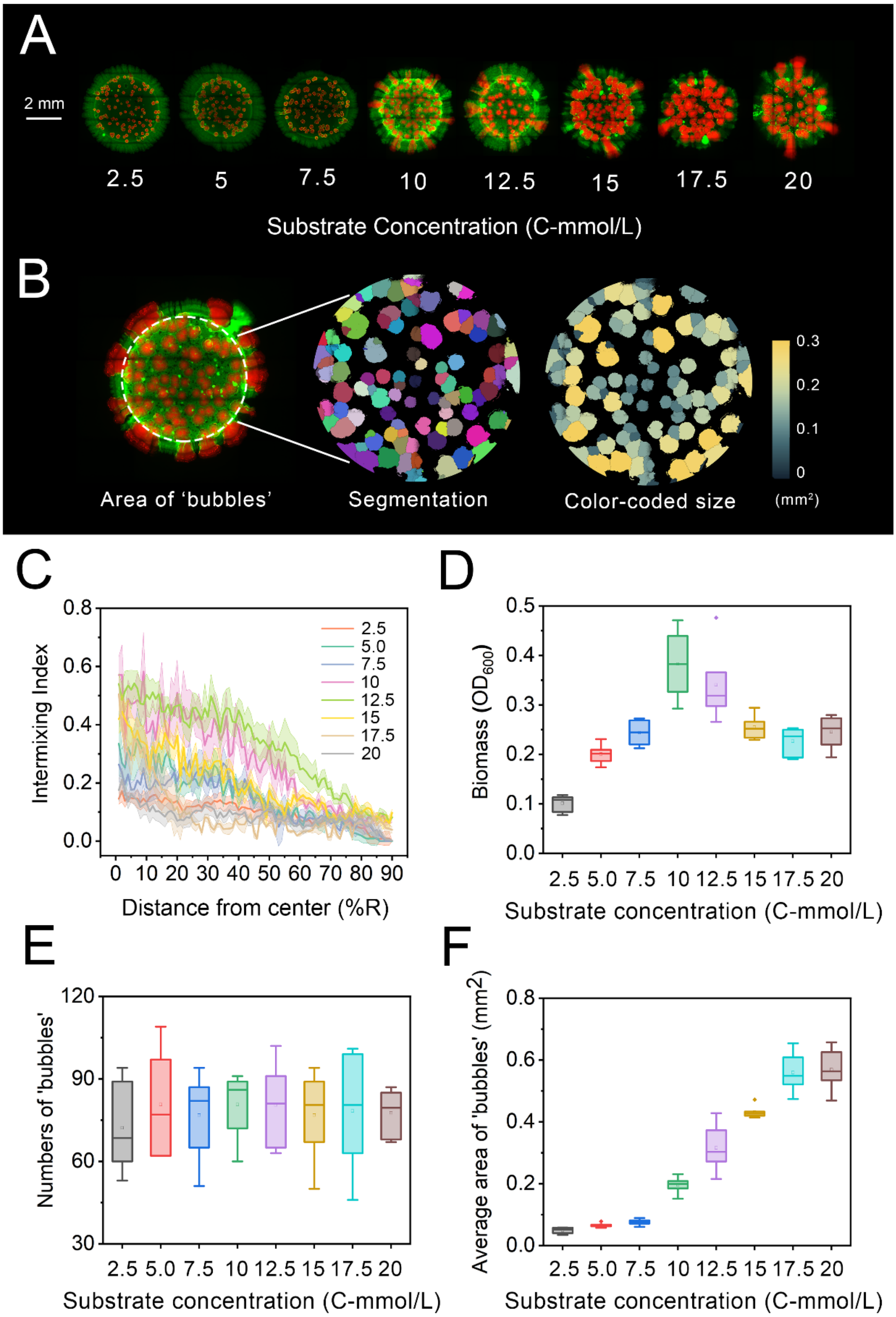


**Figure S2** Shifts of substrate concentration does not vanish the 'bubble-jet' pattern but affect the geometry of the 'bubble' structures inside the patterns. (A) Representative colony patterns from the pattern formation assays of our synthetic salicylate-degrading community across eight different initial substrate concentrations. These images clearly showed that the basic morphology of 'bubble-jet' pattern developed independent of substrate concentration. (B) Workflow of the image analysis of the 'bubble' area. 'Bubbles' formed by AN0010 cells were segmented and analyzed to get its area size ($mm^2$). In the right graph, bubbles are color-coded based on their individual area size, with brighter colors indicating larger sizes. Details of this analysis can be found in Supplementary Information S1.5. (C) Analyses of intermixing index of these patterns. (D) Analyses of the biomass of these colonies. (E) Average number of the bubbles in the colony formed by SMC-mdol in different initial substrate concentration. (F) Average area size of the bubbles inside the colony formed by SMC-mdol in different initial substrate concentration. The results in (B-F) indicates that substrate concentration did affect the details of colony pattern. For example, it affects the intermixing level (B) and community productivity (C). Importantly, with an increase in substrate concentration, the number of the bubbles remained largely unchanged (E), but the average size of bubbles inside a colony significantly increased, varying from 0.0492 ± 0.010 $mm^2$ to 0.569 ± 0.0671 $mm^2$ (F).

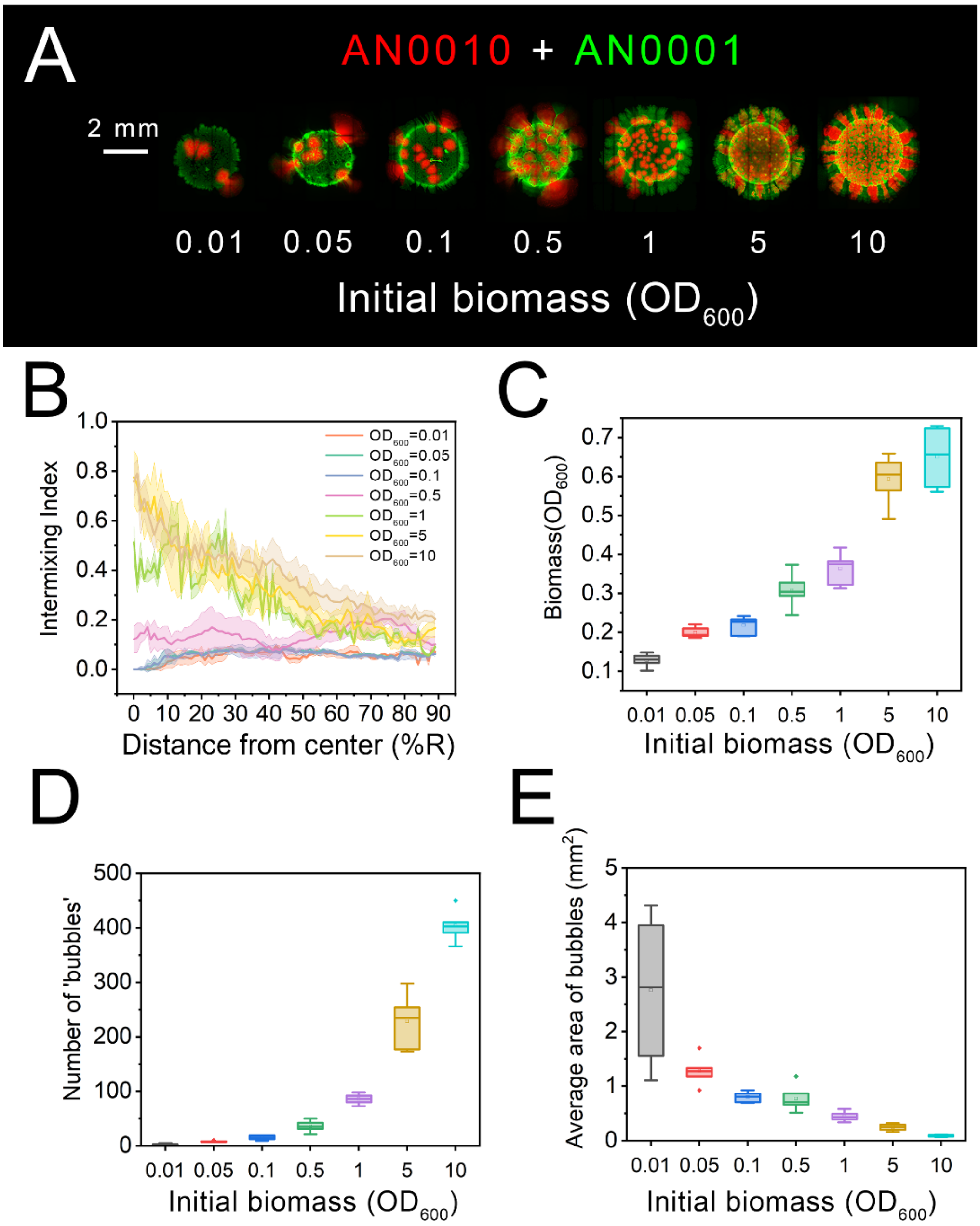


**Figure S3** Change of the density of founder cells does not vanish the 'bubble-jet' pattern but affect the geometry of the 'bubble' structures. (A) Representative colony patterns from the pattern formation assays of our synthetic salicylate-degrading community across seven different initial cell density. These images showed that the change of initial cell density did not vanish the basic morphology of 'bubble-jet' pattern. However, it did affect the details of colony pattern. For example, the intermixing level (B) and community productivity (C) increased with the initial cell density, as reported previously (9). Moreover, while the number of the bubbles rose with the increase of the initial cell density, varying from 2.16 ± 1.60 to 404 ± 27.6 (D), the bubble size significantly decreased from 2.76 ± 1.41 $mm^2$ to 0.0855 ± 0.0174 $mm^2$ (E).

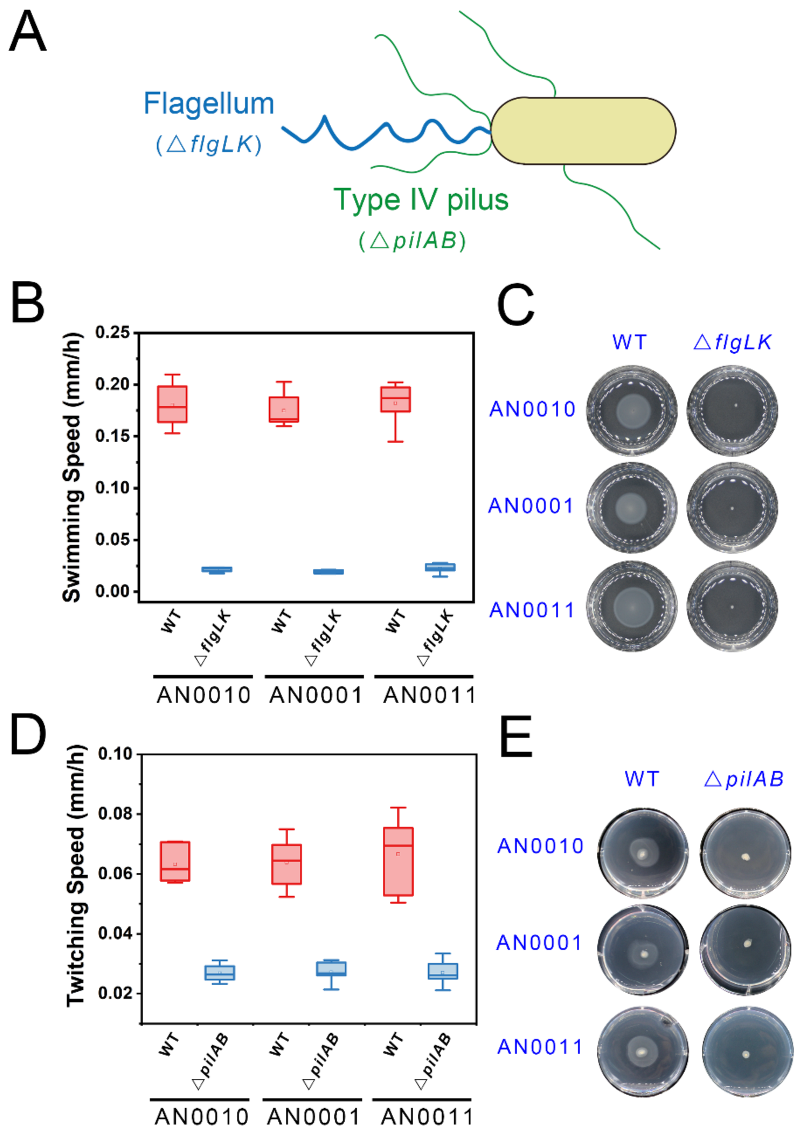


**Figure S4** Genetic manipulation to deactivate the cell appendages of our *Pseudomonas* strains. (A) Diagram indicates the appendages and the associated encoding genes considered in this study. (B-C) Swimming motility of flagellum-deficient mutants in agarose-based 'swimming' assays. (B) The swimming speed of different strains after 2-day incubation in 6-well plate. (C) Typical images of the 'swimming rings' of different strains formed inside the agarose plates. (D-E) Twitching motility of pilus-deficient mutants in agarose-based 'stab' assays. (D) The twitching speed of different strains after 3-day incubation in 6-well plate. (E) Typical images of the 'twitching rings' (faint rings that form around colonies) of different strains formed at the bottom of plates, between the plastic and the agarose. These assays were performed following the standard protocols (4), using minimum medium containing 0.3% (Swimming) or 1.5% (Twitching) agarose and 57 C-mmol/L pyruvate. The speed was determined from the analyses of 6 replicated experiments per strain, and was calculated by dividing the radius of immigration area with the incubation time.

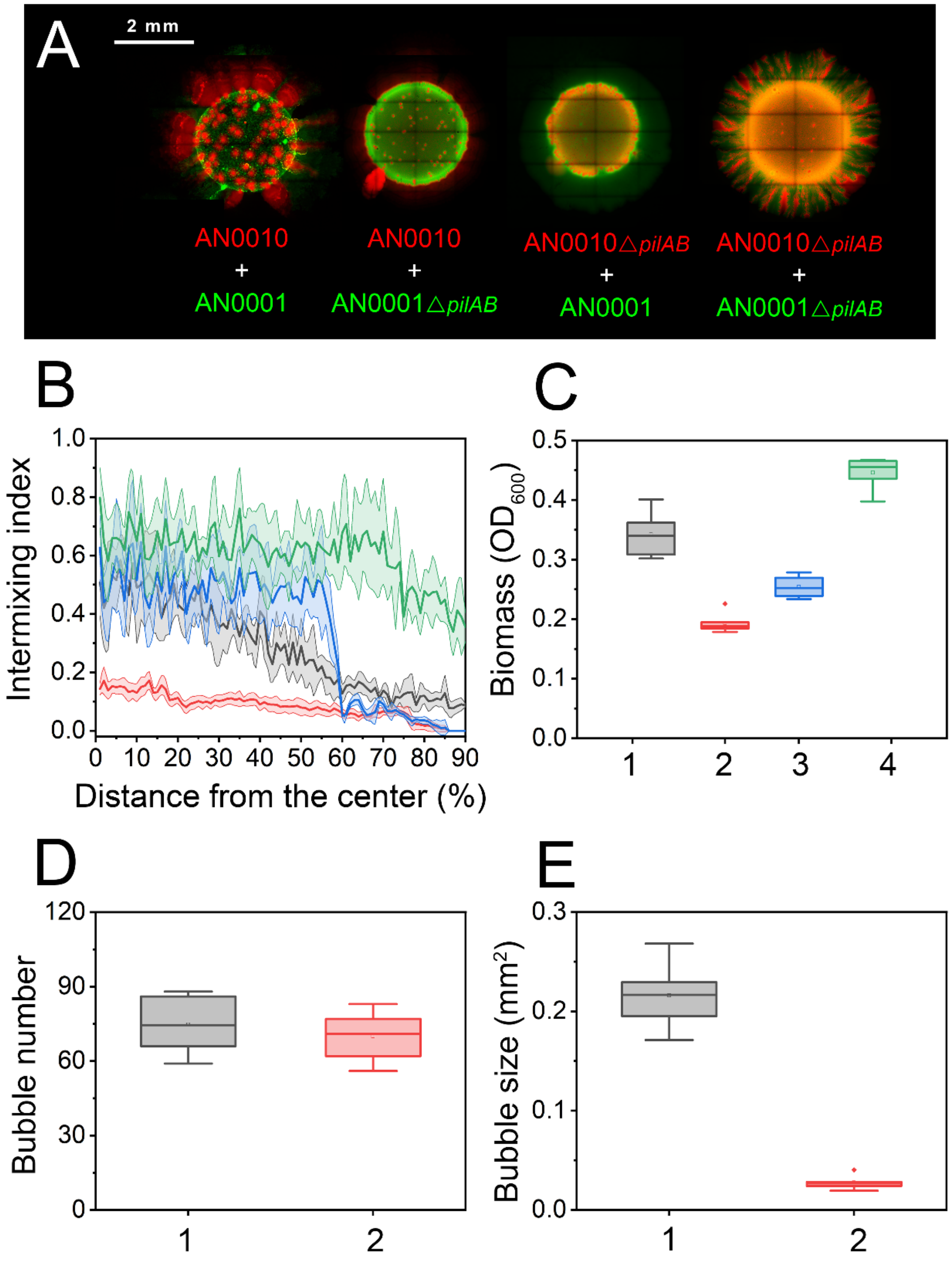


**Figure S5** Spatial patterns formed by four different combinations of the original type and the Δpili mutant of our engineered strains. (A) Representative colony patterns from the pattern formation assays of our synthetic salicylate-degrading communities composed of four different combinations of the original type and the Δpili mutant of our engineered strains. (B-C) Quantitative analyses shows the deactivation of the pili of each strain affect the intermixing level (B) and community productivity (C) of the formed patterns. (D-E) comparing with the pattern formed by the two wild type strains, in the pattern formed by the pili mutant of AN0001 and strain AN0010, while the number of the bubbles remains unchanged (D; 70.0 ± 9.9 VS 74.7 ± 11.4, unpaired two-tailed Student's t-test, $p = 0.46$), the bubble size significantly decreased (E; 0.0276 ± 0.0070 $mm^2$ VS 0.216 ± 0.0328 $mm^2$, unpaired two-tailed Student's t-test, $p = 7.83e\text{-}8$). Annotations and abbreviation in (B-E): Gray (Annotation "1"), synthetic consortium composed of strains AN0010 and AN0001; Red (Annotation "2"), synthetic consortium composed of pili mutant of strain AN0001 and strain AN0010; Blue (Annotation "3"), synthetic consortium composed of pili mutant of strain AN0010 and strain AN0001; Green (Annotation "4"), synthetic consortium composed of pili mutants of both strains AN0010 and AN0001.

Table S1 Variations when culture the two synthetic consortia in liquid medium and agarose surface initialized with different strain ratios

| | Liquid cultivation | | Surface cultivation | |
|---|---|---|---|---|
| | Frequency of AN0010 | Biomass ($OD_{600}$) | Frequency of AN0010 | Biomass ($OD_{600}$) |
| Wide-type community | 0.0236 | 0.00288 | 0.0304 | 0.00734 |
| Pili-mutant community | 0.0155 | 0.00360 | 0.00181 | 0.00237 |
